# Matrix-assisted laser desorption ionization time-of-flight mass spectrometry identification of methanogens of clinical interest

**DOI:** 10.1101/2021.11.23.469664

**Authors:** Cheick Oumar Guindo, Lynda Amir, Carine Couderc, Michel Drancourt, Ghiles Grine

## Abstract

Methanogens, the archaea uniquely detoxifying fermentative hydrogen into methane in the digestive tract, are increasingly detected in pathology situations, rendering their rapid identification mandatory. We improved the experimental protocol to identify broth-cultured methanogens by matrix-assisted laser desorption time-of-flight mass spectrometry (MALDI-TOF-MS). A database incorporating 34 reference spectra derived from 16 methanogen reference strains representative of eight species, supported further identification of 21 *Methanobrevibacter smithii* and 14 *Methanobrevibacter oralis* isolates broth-cultured from human stool and oral fluid, respectively, with scores > 2. In addition, MALDI-TOF-MS differentiated five *Methanobrevibacter smithii* genotypes incorporated in the study. Data here reported found MALDI-TOF-MS as a first line identification method for methanogens recovered from microbiota and clinical samples.

## 1. Introduction

Methanogens are acknowledged members of the digestive tract microbiota where these strictly aero intolerant archaea are detoxifying molecular hydrogen issued from anaerobic bacterial fermentation, into methane, being the only currently known sources of biotic methane [1,2]. Accordingly, the oral and gut cavities of virtually all the individuals are harboring methanogens, with respect to this indispensable physiological role [3,4]. eight different species of methanogens have been detected and cultured from oral and gut microbiota. More specifically, *Methanobrevibacter smithii* (*M. smithii*), *Methanobrevibacter oralis* (*M. oralis*) and *Methanobrevibacter massiliense* have been isolated from the oral cavity, whereas *M. smithii, M. oralis*, *Methanosphaera stadtmanae* (*M. stadtmanae*)*, Methanomassiliicoccus luminyensis* (*M. luminyensis*), *Methanobrevibacter arboriphilicus* (*M. arboriphilicus*)*, Ca*. Methanomethylophilus alvus and *Ca*. Methanomassiliicoccus intestinalis have been isolated from stools [3,5,6]. Other methanogens including *Methanosarcina mazei* (*M. mazei*), *Methanoculleus chikugoensis*, *Methanoculleus bourgensis*, *Methanobacterium congolense* have been PCR-detected in the gut microbiota but not cultured yet from clinical specimens [4].

Methanogens are emerging pathogens increasingly detected in various situations of pathology, such as dysbiosis, abscesses [2]; and more recently archaeamia [6]. In all these pathological situations, methanogens have been always detected with bacteria, including enterobacteriaceae, staphylococci and streptococci.

Owing to this increasing role in diverse pathological processes supported by the isolation and culture of methanogens in the clinical microbiology laboratory, there is a demand for a routine and specific method of identification of these still unusual microorganisms in clinical microbiology. Variable microscopic morphological aspects ranging from single coccobacillary for *M. smithii*, pairs or short chains oval rods for *M. oralis*, single cell cocci for *M. luminyensis* and pairs or tetrads cocci for *M. stadtmanae*, may not be specific enough for the accurate identification of methanogens [5]. Fluorescent *in situ* hybridization (FISH) obviously added to specificity of the microscopic examination of methanogens in clinical specimens but is yet too laborious for a routine usage [7]. Accordingly, the firm identification of cultured methanogens still relies upon the detection of species-species DNA sequences [7,8]. However, DNA-based identification methods delay identification as current archaeal DNA extraction protocols add 3- 24 hours to one-hour real-time PCR protocol or 3-hour PCR-sequencing protocol when looking for new methanogen species [9]. Potential carry-over of DNA resulting in false-positive results as well as the cost of PCR-based protocols may also limit their use for the routine identification of methanogen colonies so that a more appropriate method is desired for this purpose.

Matrix-assisted laser desorption ionization time-of-flight mass spectrometry (MALDI-TOF-MS) is currently the first-line method for identification of bacteria and yeasts cultured in clinical microbiology as this method precisely overcome the potential limitations of PCR-based methods as reported above [10]. However, this method has not been developed for the routine identification of methanogens of clinical interest since only one report has been made at a time when isolation and culture of methanogens was still in its infancy in clinical microbiology laboratories [11]. Moreover, this proof-of-concept that MALDI-TOF-MS had the potential to identify a few methanogen species was obtained from a tedious protein extraction protocol including the use of glass beads during the lysis phase [11].

Here, following improvements in experimental and informatic protocols, we are reporting on the MALDI-TOF-MS identification of broth-cultured methanogens, renewing interest in MALDI-TOF-MS-based identification of methanogens of clinical interest in the perspective of translating this approach in the routine activity of clinical microbiology laboratories.

## 2. Materials and methods

### 2.1 Methanogen clinical isolates and strains

Sixteen reference strains of methanogens available in the Collection de Souches de l’Unité des Rickettsies (CSUR, Marseille, France) were used to create a MALDI-TOF-MS reference database. They included *M. smithii* CSURP5816 of MST genotype 3, *M. smithii* CSURP5922 of MST genotype 2, *M. smithii* CSURQ5493 of MST genotype 5, *M. smithii* CSURQ5497 of MST genotype 1, *M. smithii* CSURQ5501 of MST genotype 6, *M. oralis* CSURP5701, *M. oralis* CSURQ5479, *M. oralis* CSURQ5481, *M. oralis* CSURQ5483, *M. oralis* CSURQ5485, *M.* s*tadtmanae* CSURP9634, *M. arboriphilicus* CSURP9635*, M. luminyensis* CSURP9636, *Methanobacterium beijingense* (*M. beijingense*) CSURP9638, *M. mazei* CSURP9637, and *Methanosarcina barkeri* (*M. barkeri*) CSURP9601. While *M. arboriphilicus* CSURP9635, *M. beijingense* CSURP9638, *M. mazei* CSURP9637, *M. stadtmanae* CSURP9634, and *M. barkeri* CSURP9601 have been primarily purchased from the German Collection of Microorganisms and Cell Cultures (DSMZ) (Braunschweig, Germany), *M. smithii*, *M. oralis* and *M. luminyensis* isolates have been made in our laboratory, respectively from human stool and oral fluid, as previously described [12–14]. These 16 reference strains of methanogens were subcultured in broth using a culture protocol previously established in our laboratory [15] and their identification was firmly confirmed by PCR-sequencing of the 16S rRNA gene, as previously described [16]. Further clinical isolates were made from mucosa-associated specimens as previously described [13,14]. In total, 35 clinical isolates included 21 stool isolates and 14 oral swab isolates, identified by PCR-sequencing targeting 16S rRNA Archaea and cultured in broth for nine days before being assessed by MALDI-TOF-MS against the MALDI-TOF reference database. Furthermore, we performed multispacer sequence typing (MST) of *M. smithii* 21 clinical isolates. The multispacer sequence typing technique and analysis was performed on each of the *M. smithii* clinical isolates according to the previously described protocol [9,17–19]. This part of the study was made in agreement with the positive advice of the IHU Méditerranée Infection Ethics Committee (n° 2016-020).

### 2.2 Methanogens MALDI-TOF-MS reference database

We used a protocol of the manufacturer MALDI Biotyper® (Bruker Daltonics, Wissembourg, France), adding two additional washing steps to remove the culture medium as well as extending the centrifugation time after the ethanol addition step. Briefly, one milliliter of archaeal suspension transferred into a sterile Eppendorf tube (Fisher Scientific, Illkirch, France) was centrifuged at 17,000 g for 30 minutes. The pellet was suspended into 500 μL of High Purity Liquid Chromatography water or HPLC water (VWR International, Strasbourg, France), vortexed, and centrifuged at 17,000 g for 10 minutes and this washing step was repeated twice. The pellet was suspended in 300 μL of HPLC water (VWR International), homogenized by pipetting, and 900 μL of ethanol absolute for HPLC Chromanorm (VWR International, Fontenay-sous-Bois, France) were added and homogenized by pipetting. After 5-min centrifugation at 17,000 g, the pellet was suspended in 50 μL of 70% formic acid and 50 μL of acetonitrile and vortexed for ten seconds. After a final 2-min centrifugation at 17000g, 1.5 μL of supernatant was deposited in one of a MALDI-TOF 96 MSP target polished steel BC ref number: 8280800 (Bruker Daltonik GmbH, Bremen, Germany) and 10 spots were deposited for each methanogen species. After drying, each spot was coated with 1.5 μL of a matrix solution consisting of saturated α-cyano-4-hydroxycynnamic acid or HCCA (C2020 Sigma, Lyon, France), 50% acetonitrile (CarboErba for HPLC) 2,5% trifluoroacetic acid or TFA (Uvasol for spectrometry, Sigma-Aldrich, Dorset, UK) and 47.5% HPLC water (VWR International). After drying in ambient air, the target plate was introduced into the Microflex LT® MALDI-TOF-MS device (Bruker Daltonics). Each spot was then analyzed with the help of FlexControl version 3.4 acquisition software and MALDI Biotyper® Compass version 4.1.80 (MBT Compass) analysis software. Positive control consisted of a protein extract Bacterial Test Standard Bruker ref: 8255343. Non-inoculated media and matrix solutions were used as negative controls. Also, to prevent any cross-contamination, every MALDI-TOF plate (Bruker Daltonics) used in the study, was thoroughly cleaned to remove any previous deposits of other microorganisms. The MALDI-TOF plate was first soaked in 70% ethanol (Absolute for HPLC Chromanorm) for 15 min then rinsed with HPLC water (VWR International). We subsequently stripped the MALDI-TOF plate with 500 μL of 80% TFA (Sigma-Aldrich) while gently scrubbing and then rinsed with HPLC water (VWR International), to thoroughly clean the plate under a chemical hood.

### 2.3 Reference MALDI-TOF profiling reproducibility

For each of the 16 archaea reference strains tested, ten deposits were made on MALDI-TOF-MS plate and all experiments were performed on day 9 of culture corresponding to the optimal growth period.

### 2.4 Blind MALDI-TOF identification and clustering of methanogen clinical isolates

MALDI Biotyper® version 4.1.80 software (Bruker Daltonics) was used to create reference spectra for blind identification and to create dendrograms.

Spectral references (MSP) are created using Biotyper® MSP Creation Standard Method. Five raw spectra were used for creating an MSP for each species. We performed the experiments in triplicate and a total of 34 MSP were found to be of very good quality and therefore used as a reference database. We compared the 35 clinical isolates to this database.

## 3. Results and discussion

We improved the MALDI-TOF-MS protocol for identifying cultured methanogens after a specific database was created, allowing for the rapid and routine identification of colonies issued from clinical specimens collected from mucosa-associated specimens and pathological specimens. The data here reported were validated by the negativity of the negative controls introduced in every one of the experimental steps and the reproducibility of data which were correlated with PCR-sequencing molecular identification used as the gold standard.

The database was created after three independent runs, each incorporating 10 spots for each one of 16 methanogen reference strains. We obtained a total of 270 spectra and 135 spectra (50%) analyzed of good quality, were incorporated in the database. We therefore set up a database with 34 reference spectra. We then blindly tested the identification of these 16 reference strains using the updated local database by analyzing five spots for each of them. We obtained a correct identification of these different spots with scores > 2 for each of the reference strains. We observed a unique protein profile for each one of methanogen reference strains studied (Fig.1 and Fig.2). There are 15 different MST genotypes of *M. smithii* among *M. smithii* strains known in humans according to a previous study [17]. The multispacer sequence typing is used as the first-line method for genotyping *M. smithii* strains found in humans. It allows the differentiation of *M. smithii* strains isolated from humans [17]. We observed a difference in the number and intensity of spectra between *M. smithii* strains from different MST genotypes (Fig.2B). These observations proved the reproducibility and the inter-species specificity of the method. These data confirmed the ability of MALDI-TOF-MS to identify archaea cultured in broth, as previously reported in a proof-of-concept study using a difficult protein extraction protocol [11]. We further tested our database against all the MALDI-TOF-MS databases available in our laboratory, incorporating 16,908 spectra representative of 12,290 different microbial species i.e., 4,335 bacteria, 5,989 parasites and 1,966 fungi, and no identification was observed which proved the specificity of our spectra and an absence of contaminating spectra. Genotyping the 21 clinical isolates of *M. smithii* revealed five different genotypes. Genotype 1 was found in 10/21 (47.76%); genotype 2 in 5/21 (23.80%); genotype 3 in 3/21 (14.28%); genotypes 5 in 2 /21 (9.52%) and genotype 6 in 1/21 (4.76%). These results confirm the predominance of the MST1 genotype in clinical isolates of *M. smithii* found in the human digestive tract [9,18].

**Figure 1.**
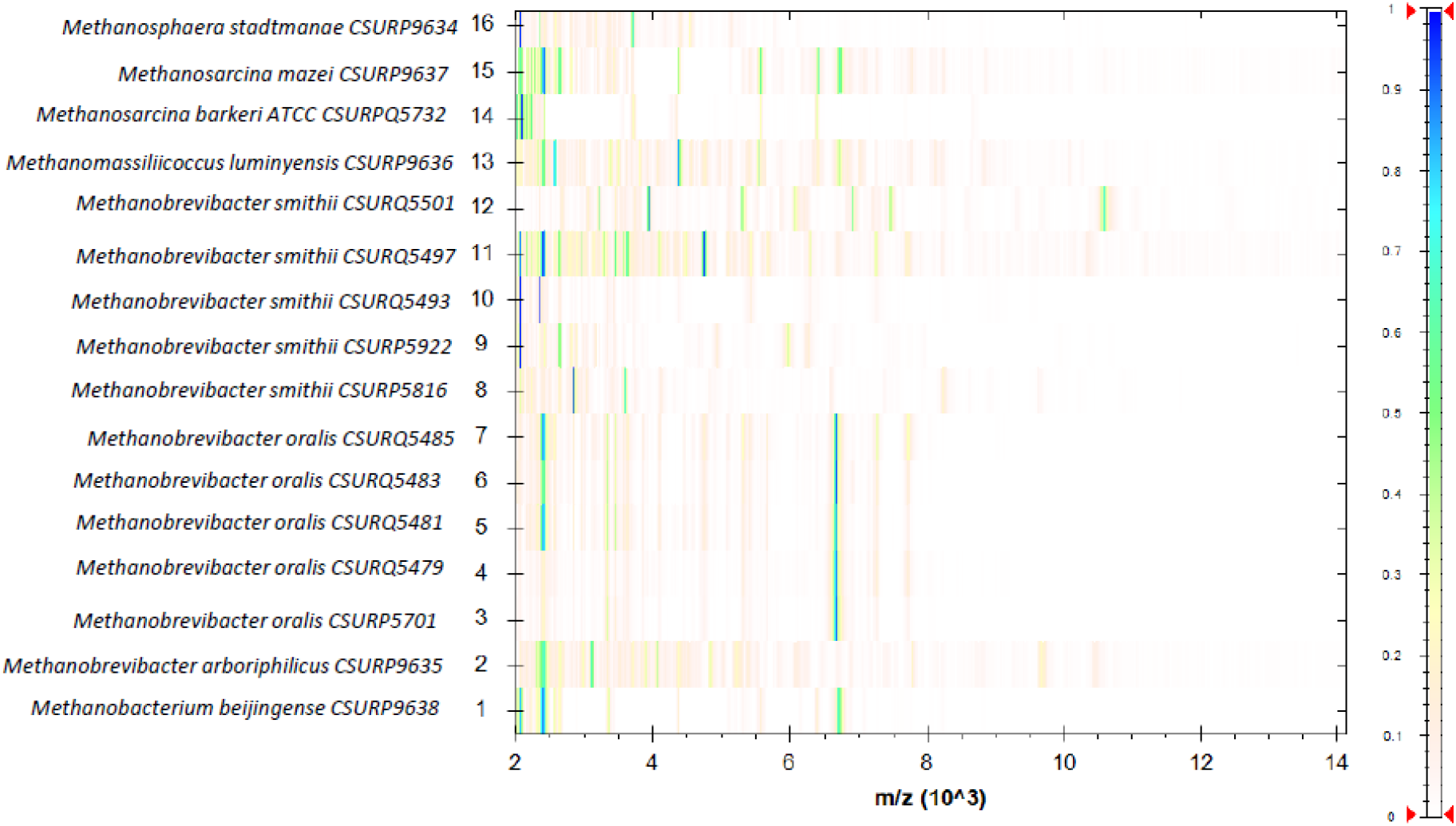
Normalized gel view of the obtained nine Archaea spectra created by MALDI Biotyper® version 4.1.80 software (Bruker Daltonics).

**Figure 2.**
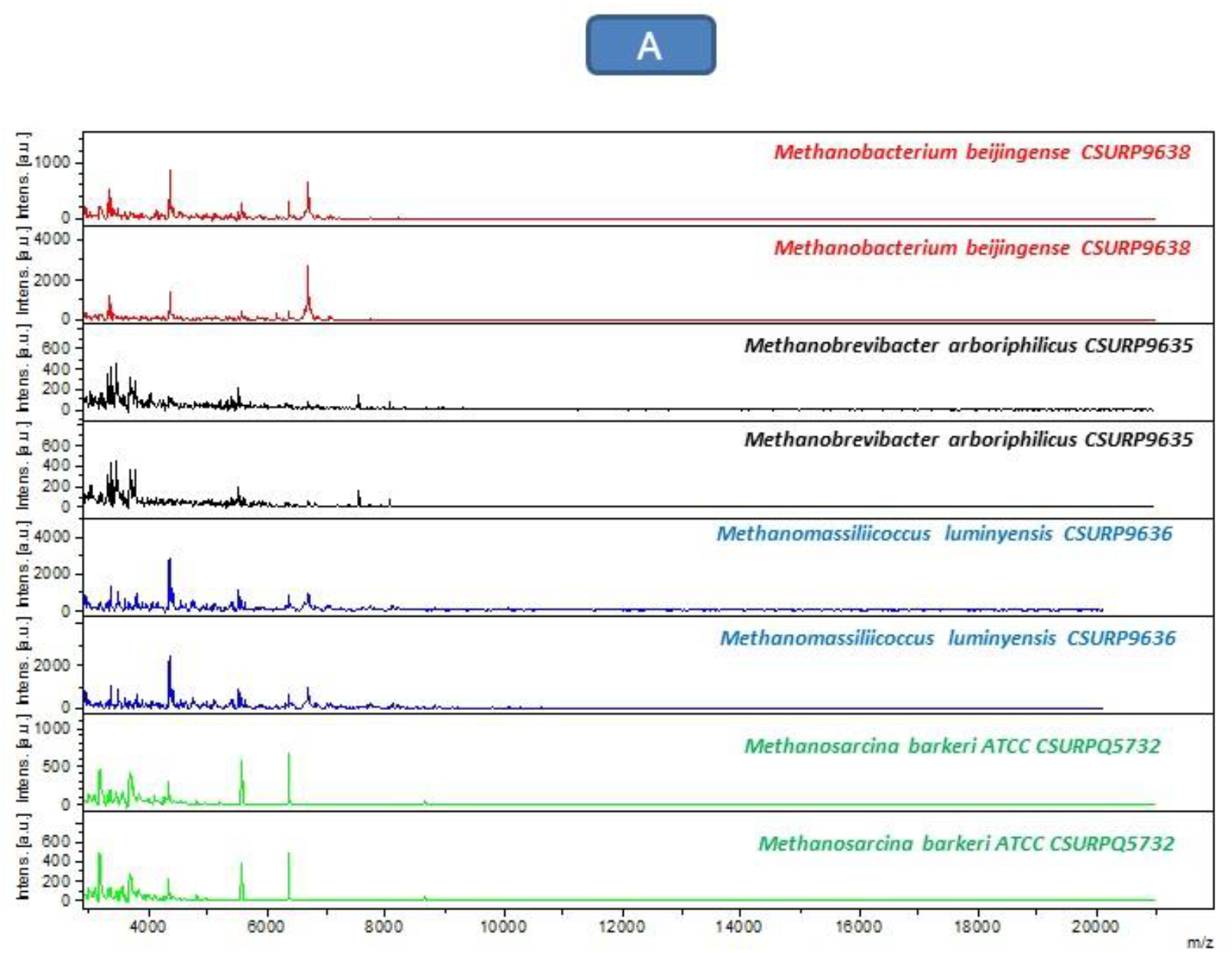
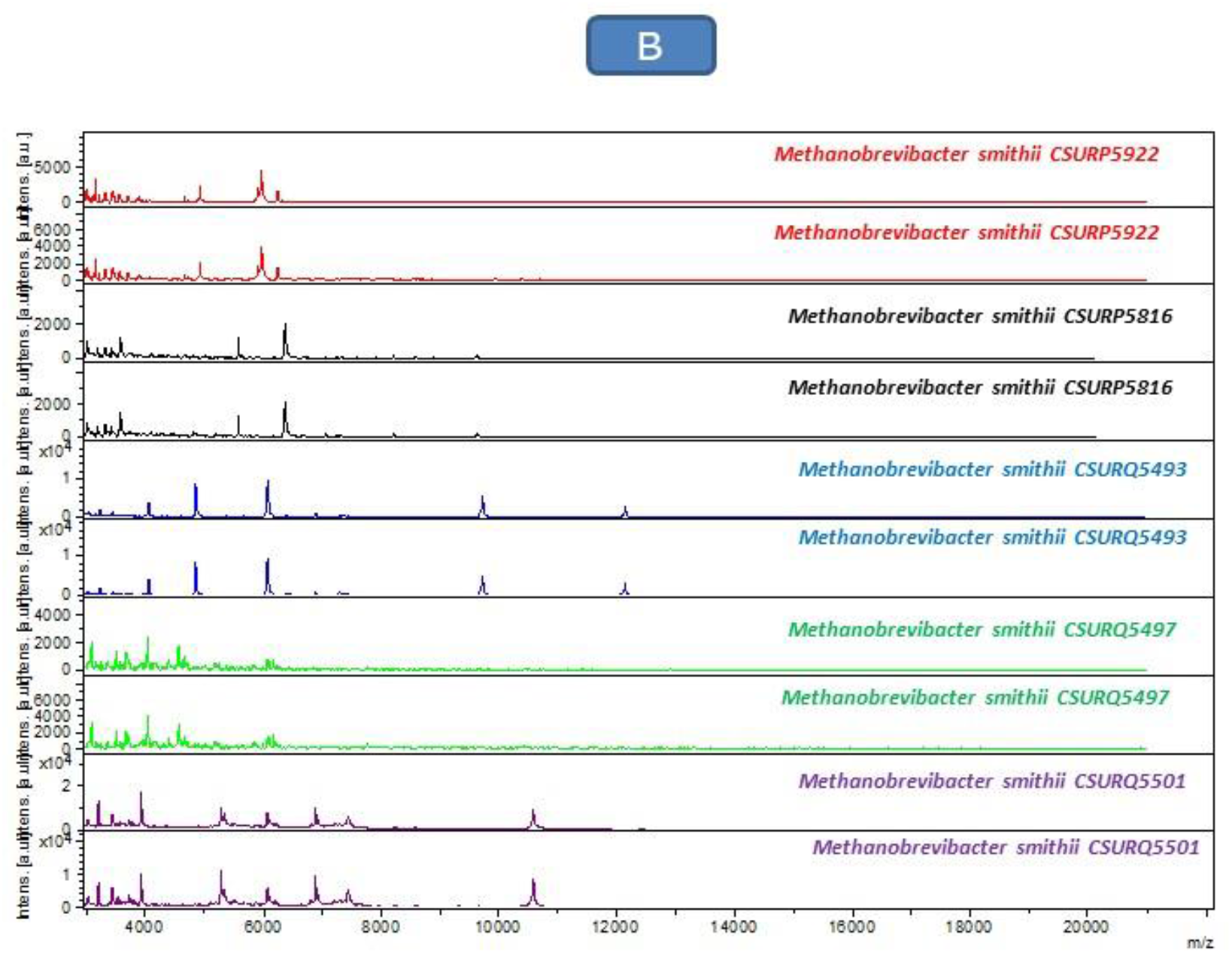
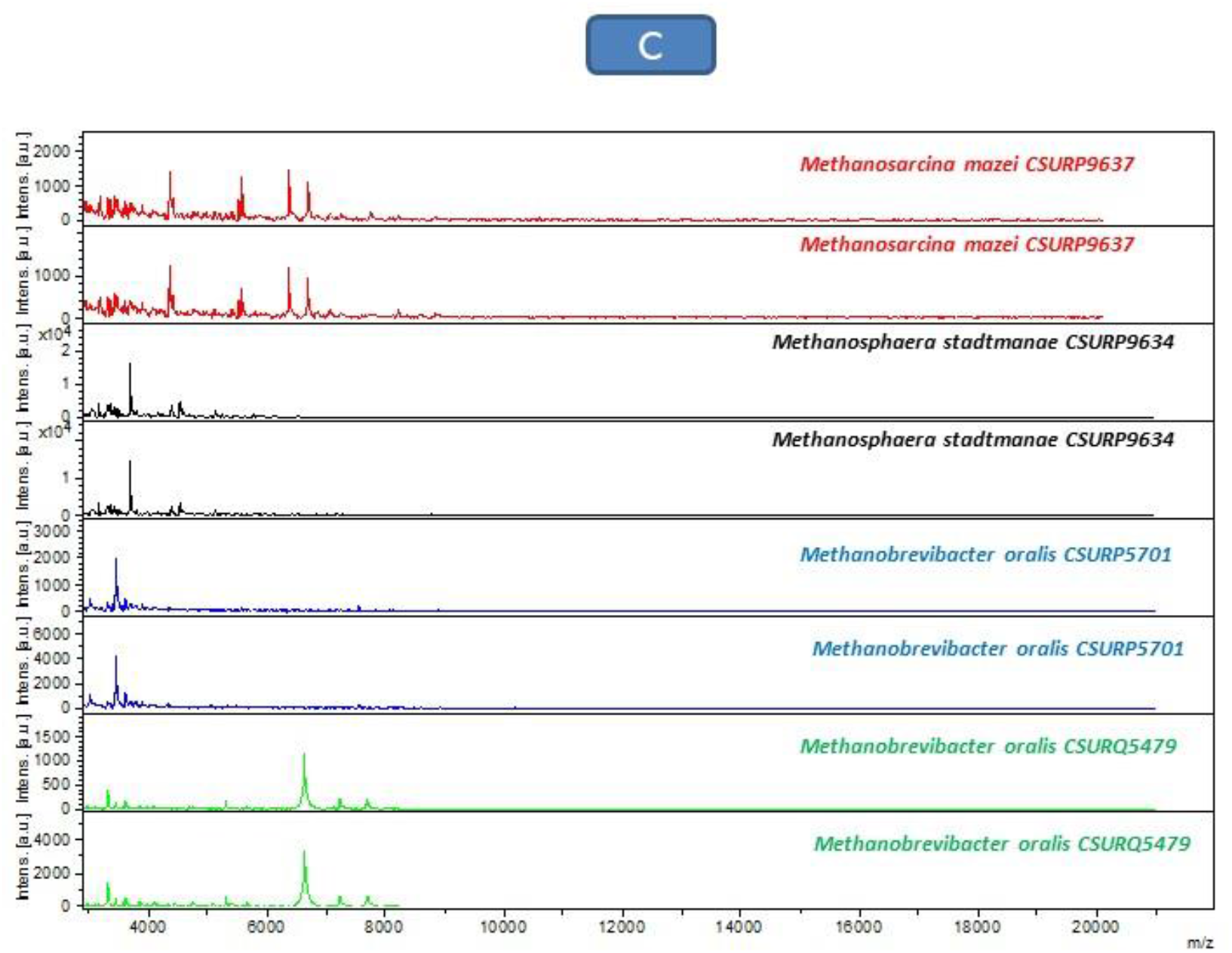
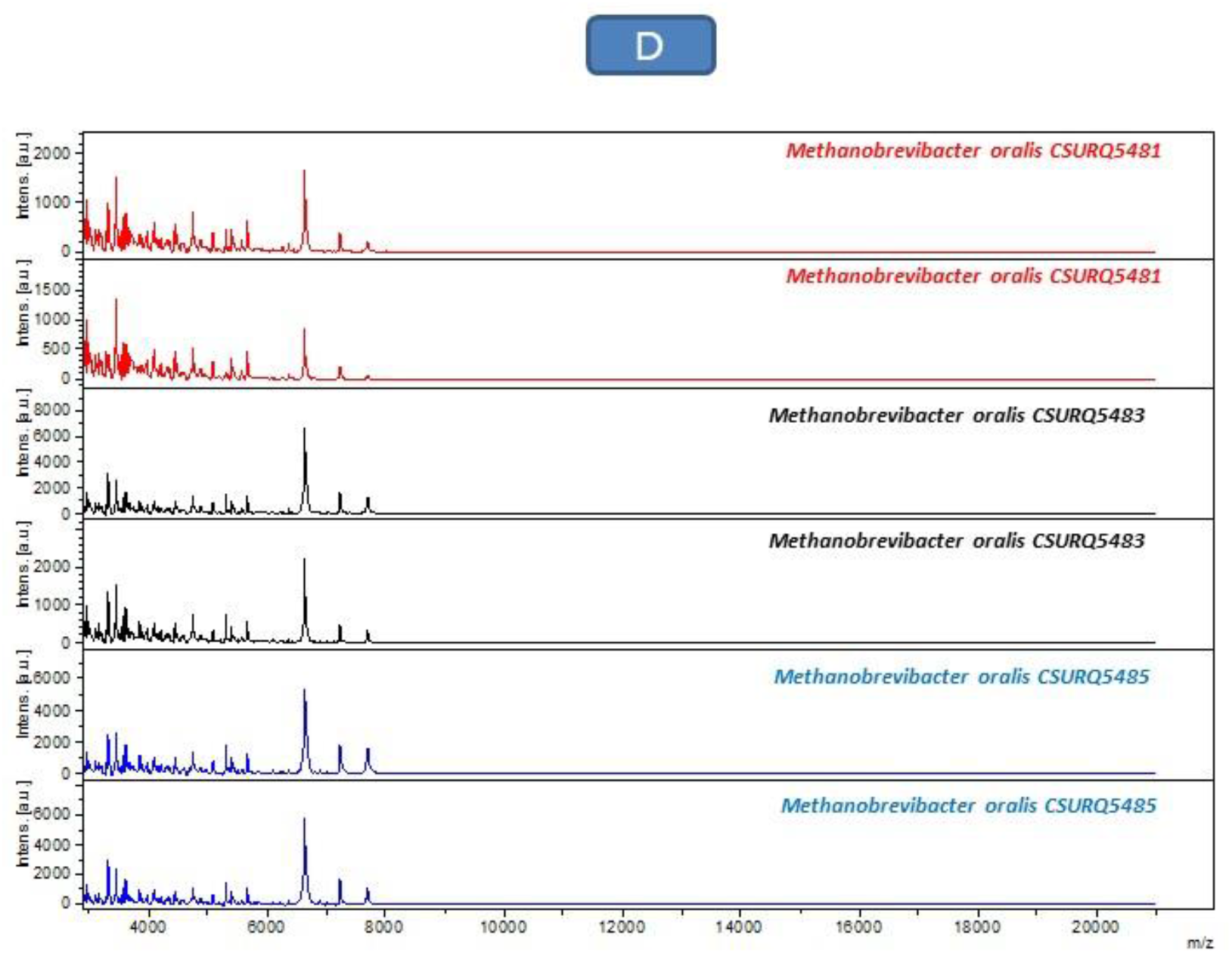
Good quality spectra of 16 methanogen reference strains studied visualized using FlexAnalysis software. (A: Spectral profiles of *M. beijingense* CSURP9638, *M. arboriphilicus* CSURP9635, *M. luminyensis* CSURP9636, *M. barkeri* CSURQ5732 reference strains / B: Spectral profiles of *M. smithii* CSURP5922, *M. smithii* CSURP5816, *M. smithii* CSURQ5493, *M. smithii* CSURQ5497, and *M. smithii* CSURQ5501 reference strains / C: Spectral profiles of *M. mazei* CSURP9637, *M. stadtmanae* CSURP9634, *M. oralis* CSURP5701, *M. oralis* CSURQ5479 reference strains / D: Spectral profiles of *M. oralis* CSURQ5481, *M. oralis* CSURQ5483, and *M. oralis* CSURQ5485 reference strains).

We then used the reference database for the identification of clinical isolates of *M. smithii* and *M. oralis* obtained by broth culture of human stool and oral fluid, respectively. We obtained a correct identification of all the clinical isolates tested with scores >2 i.e., 21/21 (100%) for *M. smithii* and 14/14 (100%) for *M. oralis,* respectively. We noticed that for a good identification by MALDI-TOF-MS of the *M. smithii* species, it is necessary to incorporate a larger set of reference spectra of *M. smithii* of different MST genotype in the reference database due to the diversity observed within this species [9,17,18], which results in different spectral profiles depending on each MST genotype as obtained in our study. To confirm this hypothesis, we repeated the blinding test with only two reference strains of *M. smithii* with different MST genotype i.e., *M. smithii* CSURP5816 of MST genotype 3, and *M. smithii* CSURP5922 of MST genotype 2. We observed that with these two reference strains, we obtained only 23% (5/21) of correct identification with scores> 2 and 19% (4/21) with scores> 1.7. Our results demonstrate the need for rapid expansion of the MALDI-TOF-MS database to incorporate isolates of clinical interest.

The results here confirmed the proof-of-concept study carried out in our laboratory in which MALDI-TOF-MS was able to perfectly identify the methanogens and environmental archaea [11]. We further observed that MALDI-TOF-MS could differentiate between different genotypes of *M. smithii*, opening the possibility to use MALDI-TOF-MS as a first line typing method for this species of prime clinical interest. We are implanting the experimental protocol and MALDI-TOF-MS methanogen database in the workflow of our clinical microbiology laboratory, for the routine, rapid identification of methanogens of clinical interest. The MALDI-TOF MS database created in this study are available on the website of the University-Hospital Institute (IHU) Méditerranée Infection on the link https://www.mediterranee-infection.com/acces-ressources/base-de-donnees/urms-data-base/, for use by the scientific community.

## Acknowledgements

We thank Amael FADLANE for his help and assistance in the conservation of methanogen strains.

## Author contributions

COG Collection of samples, study design, manipulations, data information, results interpretation, draft writing; LA Study design, data information; CC Results interpretation, draft writing; MD Supervision, study design, draft writing; GG Supervision, study design, draft writing.

